# Opposing roles of dopamine receptor D1- and D2-expressing neurons in the anteromedial olfactory tubercle in acquisition of place preference in mice

**DOI:** 10.1101/421602

**Authors:** Koshi Murata, Tomoki Kinoshita, Yugo Fukazawa, Kenta Kobayashi, Akihiro Yamanaka, Takatoshi Hikida, Hiroyuki Manabe, Masahiro Yamaguchi

## Abstract

Olfaction induces adaptive motivated behaviors. Odors associated with food induce attractive behavior, whereas those associated with dangers induce aversive behavior. We previously reported that learned odor-induced attractive and aversive behaviors accompany activation of the olfactory tubercle (OT) in a domainand cell type-specific manner. Odor cues associated with a sugar reward induced attractive behavior and c-fos expression in the dopamine receptor D1-expressing neurons (D1 neurons) in the anteromedial OT. In contrast, odor cues associated with electrical shock induced aversive behavior and c-fos expression in the D2 neurons in the anteromedial OT, as well as the D1 neurons in the lateral OT. Here, we investigated whether the D1 and D2 neurons in the anteromedial OT play distinct roles in attractive or aversive behaviors, using optogenetic stimulation and real-time place preference (RTPP) tests. Mice expressing ChETA (ChR2/E123T)-EYFP in the D1 neurons in the anteromedial OT spent a longer time in the photo-stimulation side of the place preference chamber than the control mice expressing EYFP. On the other hand, upon optogenetic stimulation of the D2 neurons in the anteromedial OT, the mice spent a shorter time in the photo-stimulation side than the control mice. Local neural activation in the anteromedial OT during the RTPP tests was confirmed by c-fos mRNA expression. These results suggest that the D1 and D2 neurons in the anteromedial OT play distinct roles in attractive and aversive behaviors, respectively.

## 1 Introduction

Odor sensation elicits various motivations, which enable adaptive behavioral responses such as obtaining food rewards or avoiding potential dangers (Doty, 1986). Although some odorants elicit innate motivated behaviors in mice, such as fear responses to predator odors(Kobayakawa et al., 2007; Saito et al., 2017) or attractive responses to social odors (Inokuchi et al., 2017), animals can acquire appropriate behaviors to odor cues according to their experience, through odor-reward or odor-danger associative learning. However, the neural circuit mechanisms engaged in these odor-induced adaptive behaviors are still unclear.

Recent studies have revealed the importance of the olfactory tubercle (OT) in the odor-induced motivated behaviors (DiBenedictis et al., 2015; Gadziola et al., 2015; Yamaguchi, 2017; Zhang et al., 2017a; Murofushi et al., 2018). The OT is a part of the olfactory cortex that receives olfactory inputs directly from the olfactory bulb as well as indirectly from other parts of the olfactory cortex and the orbitofrontal cortex (Shepherd, 2004; Zhang et al., 2017b). The OT is also a part of the ventral striatum, in addition to the nucleus accumbens (NAc), that receives massive dopaminergic inputs from the ventral tegmental area (Ikemoto, 2007; Zhang et al., 2017b; Poulin et al., 2018). The OT is composed of three major types of neurons: medium spiny neurons, dwarf cells, and granule cells (Millhouse and Heimer, 1984; Xiong and Wesson, 2016). The medium spiny neurons are distributed in the whole OT, forming the layer II (dense cell layer) of the cortex-like region (Millhouse and Heimer, 1984). A majority of the medium spiny neurons in the OT as well as the NAc and dorsal striatum express either dopamine receptor D1 or D2 (Yung et al., 1995; Murata et al., 2015). Dwarf cells are clustered in the lateral part of the OT, forming the cap region, which is interspersed throughout the antero-posterior axis (Hosoya and Hirata, 1974; Murata et al., 2015). The dwarf cells are considered a smaller type of the medium spiny neurons, and express D1 but not D2 (Murata et al., 2015). Granule cells are clustered through the anteromedial surface to the central deep part of the OT, forming the Islands of Calleja, which is presumably a continuous structure (Fallon et al., 1978; de Vente et al., 2001). The granule cells weakly express D1, and do not express D2 (Murata et al., 2015). In addition to these three types of neurons in the striatal component, the OT contains the ventral pallidal component and axon bundles that project from the striato-pallidal structure to other brain areas, forming the medial forebrain bundle (Heimer, 1978).

In our previous study, we divided the OT into domains, using the cap and Islands of Calleja as a landmark, and mapped c-fos expression when mice showed learned odor-induced attractive or aversive behaviors (Murata et al., 2015). Odor cues associated with a sugar reward induced attractive behavior and c-fos expression in the D1-expressing neurons (D1 neurons) in the cortex-like region of the anteromedial domain, which is covered by the superficially located Islands of Calleja. In contrast, odor cues associated with electrical shock induced aversive behavior and c-fos expression in the D2-expressing neurons (D2 neurons) in the cortex-like region of the anteromedial domain, as well as D1 neurons in the cap and cortex-like regions of the lateral domain, which is surrounded by the cap region. These results raise the possibility that the D1 and D2 neurons in the anteromedial OT play opposing roles in odor-guided motivated behaviors. Consistent with this idea, the D1 and D2 neurons in the NAc have distinct roles in attractive and aversive learning (Hikida et al., 2010). Here, we investigated whether activation of the D1 and D2 neurons induces attractive and aversive behaviors, respectively, by combining optogenetic stimulation and real-time place preference (RTPP) tests (Zhang et al., 2017a).

## 2 Materials and Methods

### Animals

All experiments were conducted in accordance with the Guidelines for Animal Experimentation in Neuroscience of the Japan Neuroscience Society, and were approved by the Experimental Animal Research Committee of University of Fukui. The D1-Cre and D2-Cre mice used were heterozygotes and bred from D1-Cre (the Mutant Mouse Resource & Research Centers, STOCK Tg(Drd1a-cre)FK150Gsat/Mmucd, stock number: 029178-UCD) (Gong et al., 2003; Gong et al., 2007) and D2-Cre (the Mutant Mouse Resource & Research Centers, B6.FVB(Cg)-Tg(Drd2-cre)ER44Gsat/Mmucd, stock number: 032108-UCD) (Gong et al., 2003; Gong et al., 2007) by mating the heterozygote transgenic mice with wild type C57BL/6J mice (Japan SLC). All mice were housed with their littermates until the surgery and then individually housed with a 12/12-h light/dark cycle. Food and water were freely available.

### Virus preparation

We used a Cre-dependent adeno-associated virus (AAV) vector encoding enhanced yellow fluorescent protein (EYFP) or ChETA(ChR2/E123T)-EYFP for cell type-specific gene delivery. We obtained AAV5-EF1a-DIO-EYFP from the UNC vector core at a titer of 3.5 × 10^12^ genome copies/mL; AAV2-EF1a-DIO-ChETA-EYFP was packaged and concentrated to a titer of 1.6 × 10^12^ genome copies/mL, as previously reported (Kobayashi et al., 2016), using the Addgene (Cambridge, MA, USA) plasmid, pAAV-Ef1a-DIO ChETA-EYFP (gift from Karl Deisseroth, # 26968 (Gunaydin et al., 2010)).

### Stereotaxic surgery

Stereotaxic surgeries were performed on mice aged 10–16 weeks. Mice were anesthetized with a mixture of three anesthetics (medetomidine, midazolam, and butorphanol) (Nakamura et al., 2017), and then placed in a stereotaxic apparatus (Narishige, SR-5M). The skull above the targeted areas was thinned using a dental drill and removed carefully. Injections were administered using a syringe pump (WPI, UltraMicroPump III) connected to a Hamilton syringe (Hamilton, RN-1701), and a mounted glass micropipette with a tip diameter of 50 μm connected by an adaptor (Hamilton, 55750-01).

We ipsilaterally injected 300 nL AAV5-EF1a-DIO-EYFP for confirmation of cell type-specific expression (Fig. 1C–D) and as a control for optogenetic stimulation, or AAV2-EF1a-DIO-ChETA-EYFP into the anteromedial OT of D1-Cre or D2-Cre mice using the following coordinates: anterior-posterior, +1.5 mm; medial-lateral, 0.7 mm from bregma; and dorso-ventral, 4.35 mm from the brain surface. Two to 3 weeks later, the mice were ipsilaterally implanted with a chronic optical fiber (numerical aperture = 0.39, 200-μm diameter; Thorlabs, CFMC12U) targeted to the anteromedial OT with the same coordinates described above. One to 2 weeks after fiber implantation, the following behavioral tests were conducted.

**Figure 1.**
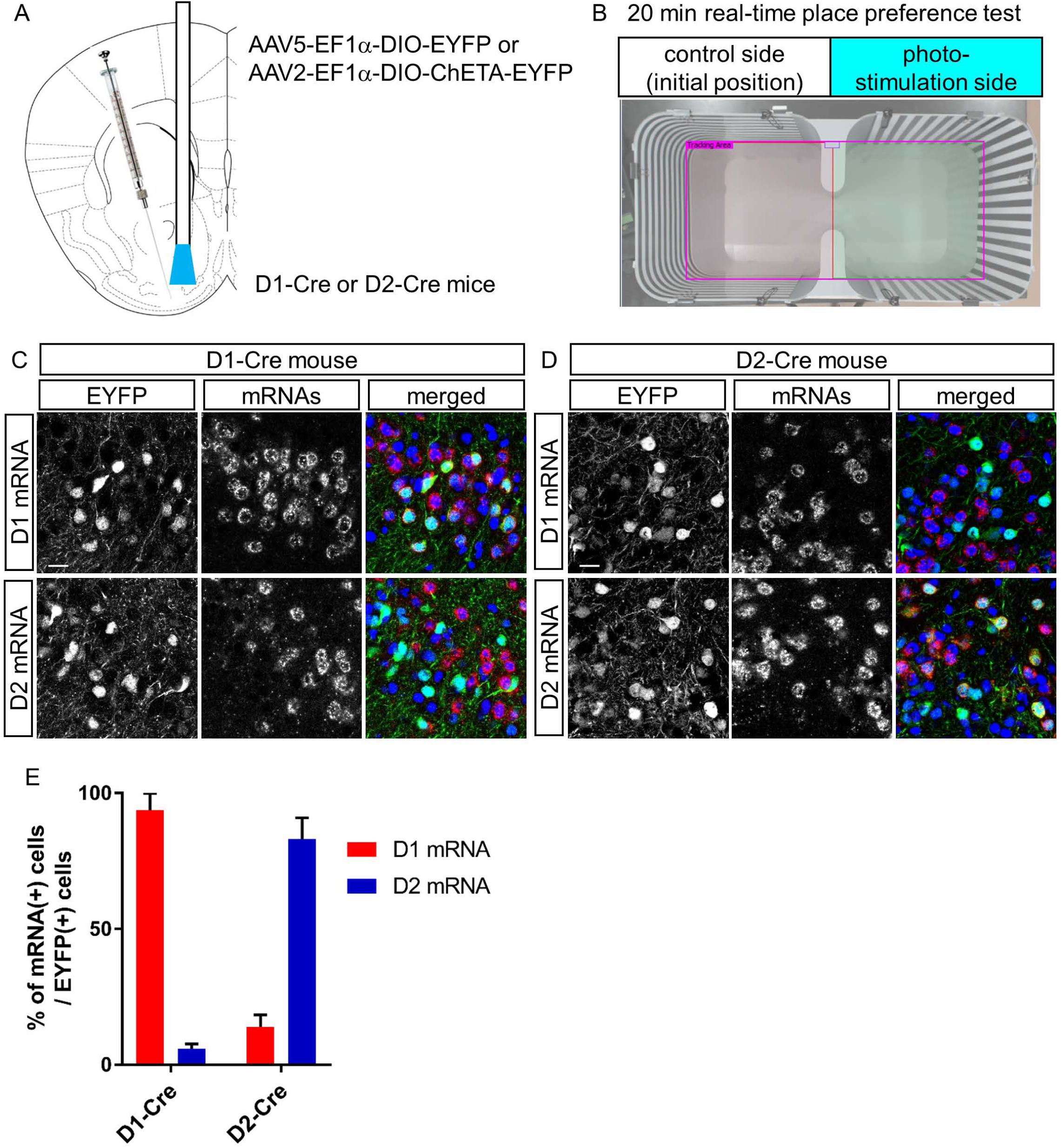
Cell type-specific gene delivery in dopamine receptor D1- and D2-expressing neurons in the anteromedial OT of the D1-Cre and D2-Cre mice using AAV vectors. (A) Schematic diagram of cell type-specific optogenetic stimulation of the anteromedial OT. We injected Cre-dependent AAVs encoding EYFP or ChETA-EYFP and implanted optic fiber canula into the anteromedial OT of D1-Cre or D2-Cre mice. (B) Place preference chamber. In the RTPP tests, mice were placed in either side, which was assigned as the control side (no photo-stimulation). Blue light was delivered when mice were in the opposite side of the initial position. (C) and (D) Confocal images of AAV-derived EYFP expressing cells (green) and D1 (upper panels) or D2 (lower panels) mRNAs (red) from D1-Cre mouse (C) or D2-Cre mouse (D). Color merged panel contains DAPI staining (blue). Scale bars: 20 μm. (E) Percentage of D1 or D2 mRNA expressing-cells among EYFP-expressing cells in D1-Cre and D2-Cre mice. Data shows mean with SD. OT, olfactory tubercle; AAV, adeno-associated virus; EYFP, enhanced yellow fluorescent protein; RTPP, real-time place preference; D1, dopamine receptor D1; D2, dopamine receptor D2; DAPI, 4′,6-diamidino-2-phenylindole; SD, standard deviation

### Optogenetic stimulation and RTPP tests

For optogenetic stimulation, the implanted optic fiber was connected to a blue light laser via patch cords with a fiber-optic rotary joint (Thorlabs, RJPSF2). All photo-stimulation experiments used 5-ms, 5–7-mW, 473-nm light pulses at 20 Hz via a solid-state laser for light delivery (Ultra Laser, CST-L-473-50-OEM) triggered by a stimulator (Bio Research Center, STO2).

After being connected to the blue light laser, the mice were placed in a place preference chamber (30 [width] × 30 [depth] × 25 [height] cm) equipped with vertical or horizontal striped wall, as shown in Fig. 1B, for 20 min. The non-stimulation control side was as assigned at the start of the experiment. Laser stimulation at 20 Hz was constantly delivered when the mice crossed to the stimulation side, and stopped when they returned back to the initial non-stimulation side. All behavioral tests were recorded using a USB digital video camera (Logicool c920r), and offline analyses of the time spent in each chamber and tracking data were performed using a videotracking software (Panlab, SMART 3.0); 30 min after the end of the RTPP tests, mice were deeply anesthetized by intraperitoneal injection of sodium pentobarbital, and then fixed for histochemical analysis.

### Histochemistry

Mice were transcardially perfused with phosphate-buffered saline (PBS), followed by 4% paraformaldehyde (PFA). After cryoprotection with sucrose solution, the brain was frozen and sliced into coronal sections with a thickness of 20 μm. The sections were rinsed in PBS and 0.1 M phosphate buffer, mounted on glass slides using a paint brush, dried overnight in a vacuum desiccator, and then stored at 4 °C until histochemistry.

To confirm cell type-specific EYFP expression, we performed double fluorescent immunolabelling for EYFP and mRNAs of D1 or D2 as follows. Digoxigenin (DIG)-labeled RNA probes were prepared using an in vitro transcription kit (Roche) according to the manufacturer’s protocol with a plasmid kindly provided by Dr. Kazuto Kobayashi (Sano et al., 2003). The dried sections were fixed in 4% PFA, digested using proteinase K (10 μg/mL) for 30 min, and post-fixed in 4% PFA. After prehybridization, the sections were incubated overnight at 65 °C with DIG-labelled RNA probes. After stringent washing, the sections were incubated in 1% blocking buffer (Roche, 11096176001) for 1 h. Primary antibody against EYFP (1:1000; Medical & Biological Laboratories) and an anti-DIG antibody conjugated with alkaline phosphatase (1:500, Roche) were included in the incubation mixture. The sections were washed three times in TNT (0.1 M Tris-HCl [pH 7.5], 0.15 M NaCl, 0.1% Tween 20) and incubated with an Alexa Fluor 488-conjugated secondary antibody (1:400; Jackson ImmunoResearch Labs) for 2 h. After three washes in TNT and one wash in Tris saline (0.1 M Tris-HCl [pH 8.0], 0.1 M NaCl, 50 mM MgCl2), alkaline phosphatase activity was detected using the HNPP Fluorescence Detection Set (Roche, 11758888001) according to the manufacturer’s instructions. The sections were incubated with the substrate three times for 30 min each, and the reaction was stopped by washing the sections in PBS. The sections were then counterstained with 4′,6-diamidino-2-phenylindole diluted in PBS (2 µg/mL) for 5 min. After washing in PBS, the sections were mounted in PermaFluor (Thermo Fisher Scientific).

For c-fos mRNA detection, we performed *in situ* hybridization using DIG-labeled antisense RNA probes. The RNA probe was prepared using an in vitro transcription kit (Roche) according to the manufacturer’s protocol with a plasmid kindly provided by Dr. Hirohide Takebayashi (Bepari et al., 2012). Hybridization and washing were performed as described above. Subsequently, the sections were blocked with 10% normal sheep serum, 1% bovine serum albumin, and 0.1% Triton X-100 in PBS. Subsequently, the sections were incubated overnight at 4 °C with alkaline phosphatase-conjugated anti-DIG antibody (1:1000, Roche). The sections were washed in TNT, followed by alkaline phosphatase buffer (100 mM NaCl, 100 mM Tris-HCl [pH 9.5], 50 mM MgCl2, 0.1% Tween 20, 5 mM levamisole). The sections were treated overnight with nitro-blue tetrazolium/5-bromo-4-chloro-3’-indolylphosphate (Roche) mixture at room temperature in a dark room for color development. Then, they were rinsed in PBS and mounted in PermaFluor (Thermo Fisher Scientific).

### Microscopy and image analysis

Sections were examined using a confocal laser microscope (Olympus, FV1200) and a bright field virtual slide system (Hamamatsu Photonics, NanoZoomer). To quantify density of c-fos mRNA-expressing cells, the area of the anteromedial domain of the OT was delineated, and the number of cells was counted using Image J (National Institutes of Health).

### Statistics

Statistical significance was tested using Prism 7 (GraphPad Software). Differences were considered statistically significant at p < 0.05.

## 3 Results

To address whether D1 and D2 neurons in the anteromedial OT play distinct roles in attractive and aversive behaviors, we used optogenetic stimulation and performed RTPP tests (Fig. 1A and B). We injected AAV2-EF1a-DIO-ChETA-EYFP into the anteromedial OT of D1-Cre and D2-Cre transgenic mice; AAV5-EF1a-DIO-EYFP was injected as a control for optogenetic stimulation (Fig. 1A). At first, we examined cell type-specificity of the Cre-mediated gene expression by the AAV vector. Three weeks after injection of AAV5-EF1a-DIO-EYFP into the anteromedial OT of the D1-Cre and D2-Cre mice, we performed double fluorescence immunolabeling of EYFP and D1 or D2 mRNAs (Fig. 1C and D). In the D1-Cre mice, 94% of the EYFP(+) neurons in the cortex-like region, which were putative medium spiny neurons, expressed D1 mRNA, and 5.9% of them expressed D2 mRNA (n = 3 mice, Fig. 1C and D). On the other hand, 14% of the EYFP(+) neurons expressed D1 mRNA, and approximately 83% of them expressed D2 mRNA in the D2-Cre mice (n = 3 mice, Fig. 1C and E). These data confirmed that these Cre transgenic mice exhibited preferential expression of Cre-dependent AAV-derived genes in the D1 and D2 neurons.

We then tested the hypothesis that optogenetic activation of the D1 and D2 neurons in the anteromedial OT may play distinct roles in eliciting attractive and aversive behaviors using RTPP tests. We activated the D1 and D2 neurons using ChETA, a type of ChR2 with faster kinetics (Gunaydin et al., 2010), which possibly enabled precise timing of stimulation when mice crossed the chambers. The RTPP tests revealed that D1-Cre mice expressing ChETA spent significantly longer time in the photo-stimulation side (64% of the 20-min RTPP tests, n = 6 mice) than the control mice (52% of the 20-min RTPP tests, n = 7 mice; unpaired *t*-test: t = 3.261, df = 11) which expressed EYFP without ChETA (Fig. 2A and B). In contrast, D2-Cre mice expressing ChETA spent significantly shorter time in the photo-stimulation side (28% of the 20-min RTPP tests, n = 6 mice) than the control mice (51% of the 20-min RTPP tests, n = 7 mice; unpaired *t*-test: t = 3.994, df = 11) (Fig. 2A and B). These data suggest that activation of the D1 and D2 neurons in the anteromedial OT elicit attractive and aversive behaviors, respectively.

**Figure 2.**
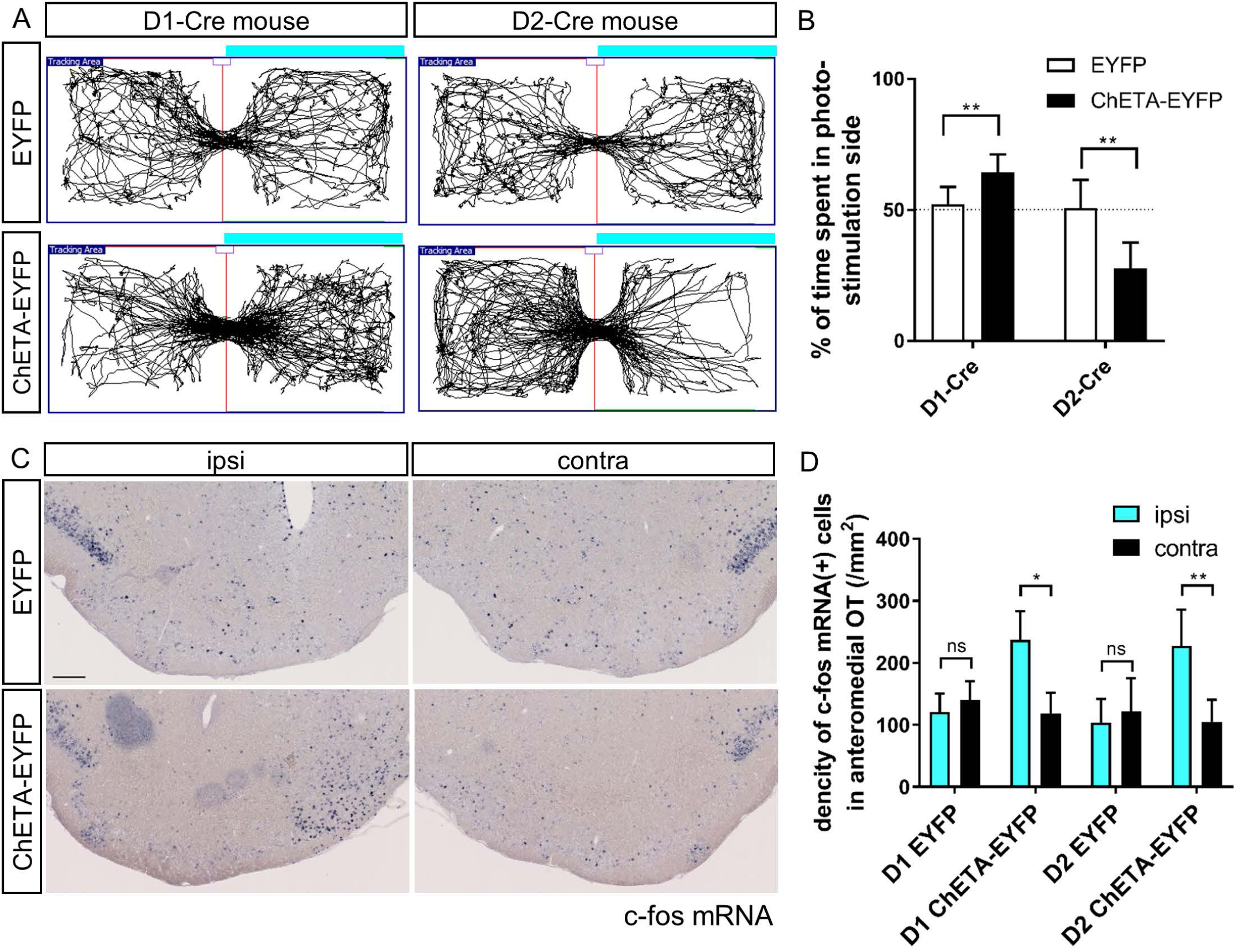
Cell type-specific effect of optogenetic stimulation of the anteromedial OT in the RTPP tests. (A) Tracking data of the RTPP tests. Right side is the photo-stimulation side. (B) Percentage of time spent in the photo-stimulation side in the 20-min RTPP tests. Data shows mean with SD. **, *p* <0.01. (C) Images of c-fos mRNA expression in the OT coronal sections after the RTPP tests. Left panels, ipsilateral; right panels, contralateral. Scale bar: 200 μm. (D) Density of c-fos mRNA-expressing cells in the anteromedial OT. Data shows mean with SD. ns, not significant; *, *p* < 0.05;**, *p* < 0.01. OT, olfactory tubercle; RTPP, real-time place preference; SD, standard deviation

After the RTPP tests, we confirmed the neural activation of the anteromedial OT by examining c-fos mRNA expression (Bepari et al., 2012). As expected, the ipsilateral side of the anteromedial OT showed a significant increase in the number of c-fos expressing cells in in both ChETA-EYFP-expressing D1-Cre (n = 5 mice; unpaired *t*-test: t = 4.628, df = 8) and D2-Cre (n = 6 mice; unpaired *t*-test: t = 4.427, df = 10) mice (Fig. 2C and D). In contrast, no clear increase in the c-fos expression in either control D1-Cre (n = 7 mice; unpaired *t*-test: t = 1.239, df = 12) or D2-Cre (n = 7 mice; unpaired *t*-test: t = 0.7561, df = 12) mice was observed (Fig. 2C and D). These results confirmed activation of the anteromedial OT by optogenetic stimulation during the RTPP tests.

## 4 Discussion

In this study, we demonstrate that cell type-specific activation of the D1 and D2 neurons in the anteromedial OT elicits attractive and aversive behaviors, respectively. To achieve selective manipulation of the D1 or D2 neurons, which are intermingled in the cortex-like region of the OT, we used D1-Cre and D2-Cre transgenic mouse lines and Cre-dependent AAV vectors (Fig. 1A and C–E). This combination enabled us to deliver AAV-derived genes preferentially to the D1 or D2 neurons in the anteromedial OT. Optogenetic activation of the D1 and D2 neurons in the anteromedial OT induced attraction to and aversion from the photo-stimulation chamber, respectively (Figs. 1B, and 2A and B). After the RTPP tests, activation of the anteromedial OT was confirmed by local increase in the c-fos expression (Fig. 2C and D). These results suggest that the D1 and D2 neurons in the anteromedial OT are involved in eliciting attractive and aversive behaviors, respectively.

Previous studies have shown that the anteromedial domain of the OT plays an important role in the reward system. Local self-administration of cocaine into the OT and NAc revealed that the anteromedial domain of the OT more robustly mediates the rewarding action of cocaine than other domains of the OT and NAc (Ikemoto, 2003). Optogenetic stimulation of the dopaminergic fiber from the ventral tegmental area to the medial OT elicits rewarding effects which generate place and odor preference (Zhang et al., 2017a). These local manipulations should exert excitatory effect on the D1 neurons via increased dopamine level in the anteromedial OT because D1 is coupled with Gs (Stoof and Kebabian, 1981). In line with these previous reports, our result directly demonstrates the role of the D1 neurons in the anteromedial OT in eliciting attractive behavior. In contrast, D2 is coupled with Gi (Stoof and Kebabian, 1981). Aversive stimuli reduce tonic firings of dopaminergic neurons (Ungless et al., 2004; Cohen et al., 2012) resulting in decreased ambient dopamine level at the target structure, which should exert excitatory effect on the D2 neurons. Consistent with the report that blunting the tonic dopamine release in the ventromedial striatum leads to conditioned place aversion (Liu et al., 2008), our results reveal the role of the D2 neurons in the anteromedial OT in eliciting aversive behavior. As we previously reported, neurons in the OT are activated by odor cues that induce motivated behaviors. The understanding of these cell type-specific roles of the D1 and D2 neurons in the anteromedial OT will provide a neural basis for odor-guided adaptive motivated behaviors.

## 5 Conflict of Interest

The authors declare that the research was conducted in the absence of any commercial or financial relationships that could be construed as a potential conflict of interest.

## 6 Author Contributions

KM designed research, performed experiments, and wrote the manuscript. TK performed experiments. YF, KK, AY, TH, HM, and MY contributed tools and reagents, and assisted in revision of the manuscript.

## 7 Funding

KM was supported by the Narishige Neuroscience Research Foundation, Takeda Science Foundation, Cosmetology Research Foundation, and JSPS KAKENHI Grant Numbers 25830032, 26120709, 16K18377, 16H01671, 17KK0190, 18H05005. Y.F. was supported by JSPS KAKENHI Grant Numbers 15H01282, 16H04662.

## 8 Acknowledgements

We thank Drs. Kazuto Kobayashi, Hirohide Takebayashi and Karl Deisseroth for their generous gift of the plasmids for RNA probes and AAV production. We also thank Hideki Yoshikawa, Eri Murai, Noriko Funabashi, and members of the Fukazawa Laboratory and Life Science Research Laboratory at University of Fukui for technical assistance.

